# Spatial memory in Alzheimer’s disease 5XFAD mice is enhanced by XPO1 inhibitor KPT-330

**DOI:** 10.1101/2024.10.21.619493

**Authors:** Shi Quan Wong, Adia Ouellette, Avery McNamara, Rachel A. Tam, Alexander Alexandrov, Acacia Nawrocik-Madrid, Jesus J. Sanchez, Brett C. Ginsburg, Arturo A. Andrade, Louis R. Lapierre

## Abstract

The proteostatic decline in Alzheimer’s disease is well established and improvement in proteostasis could potentially delay cognitive impairment. One emerging entry point to modulate proteostasis is the regulation of nucleo-cytoplasmic partitioning of proteins across the nuclear pore via karyopherins. The nuclear exportin XPO1 is a key regulator of proteostasis by driving the assembly of ribosomes and by modulating the process of autophagy. We recently found that XPO1 inhibitor KPT-330 (Selinexor), an FDA approved drug against multiple myelomas, enhances proteostasis, leading to benefits in models of neurodegenerative diseases in *C. elegans* and *Drosophila*. Here, we find that KPT-330 increases autophagy in murine neuronal cells and improves spatial memory performance in a murine model of Alzheimer’s disease (5XFAD). Unexpectedly, general amyloid deposition in several brain regions was significantly increased by KPT-330, but specific regions, especially the thalamus, displayed significantly lower deposition, suggesting that XPO1 inhibition has regional-specific effects on proteostasis and amyloid plaque formation. Altogether, we conclude that XPO1 inhibition can improve cognition via spatially-specific reductions in amyloid deposition.

## INTRODUCTION

Alzheimer’s disease (AD) remains a formidable challenge for biomedical researchers with successful therapeutics currently restricted to antibodies targeting amyloid proteins [1] with debatable effectiveness and economic benefits [2]. At its core, AD is a progressive proteinopathy that burdens neuronal function and hinders survival to the point that cognition and memory are inevitably lost. As neurons are largely post-mitotic, cells manage proteome stability via proteostatic mechanisms that primarily relies on properly controlling protein synthetic, folding and degradation machineries in order to maintain protein solubility and function. Post-mitotic systems such as the nematode *C. elegans* provide a useful model to study the biology of aging and the regulation of proteostasis. The vast majority of genetic models of enhanced proteostasis and longevity require active autophagy [3] and alterations in ribosomal assembly via nucleolar attrition [4]. Therefore, finding strategies that modulate these two key nodes of proteostasis may address the progressive proteostatic decline found in AD.

The nuclear export protein XPO1 (Exportin1, CRM1) is estimated to actively transport over 200 different proteins [5] containing nuclear export sequences (NES) across the nuclear pore, including tumor-suppressing proteins [6-8]. As such, slow reversible XPO1 inhibitors were developed to target the cysteine 528 in the NES-binding site of XPO1 [9, 10] and reduce the export of tumor-suppressing proteins, consequently preventing the loss of tumor-suppressing protein activity and reducing cancer cell proliferation [11]. Recently, the XPO1 inhibitor KPT-330 (Selinexor, XPOVIO) was approved against relapsed multiple myelomas [12]. XPO1 modulates the activity of the autophagy-lysosome pathway by controlling the localization of TFEB [8, 13]. Transcriptional activation of TFEB via XPO1 inhibition is accompanied by increases in lifespan in wild-type *C. elegans* and *Drosophila* afflicted with an ALS-related mutation. Loss of XPO1 activity also reduces overall rRNA levels, decreases ribosomal assembly, downregulates transcriptionally and translationally Fibrillarin, and restricts nucleolar size via autophagy protein GABARAP [14]. XPO1 inhibition therefore represents an emerging strategy to maintain proteome stability and function by leveraging different regulatory nodes of proteostasis.

In this study, we tested KPT-330 in cultured murine neuroblasts as well as a murine model of Alzheimer’s disease in order to evaluate its effect on the proteostatic decline accompanied by the progressive amyloid deposition, and its potential in improving cognitive functions. We find that KPT-330 prevents the loss of cognition in 5xFAD mice via region-specific reductions in amyloid deposition, suggesting that nuclear export inhibition can preserve key functionalities during the development of Alzheimer’s disease.

## RESULTS

### KPT-330 enhances autophagy in murine neuroblastoma Neuro2a cells

Reducing the activity or the expression of XPO1 was previously demonstrated to enhance proteostasis, autophagy and lifespan in model organisms [8, 13]. Therefore, we sought to evaluate the impact of the loss of XPO1 activity on the onset of Alzheimer’s disease, a neurodegenerative disease characterized by autophagic dysfunction and proteostatic collapse. To reduce XPO1 activity, we opted to use KPT-330 (Selinexor), an FDA-approved XPO1 inhibitor [15] of relatively small size (443 Da) that can cross the blood brain barrier. To assess the impact of XPO1 inhibition on autophagic degradation, we initially tested the effect of silencing *XPO1* in murine neuroblastoma Neuro2a cells. *XPO1 silencing* led to significant reductions in both its transcript and protein levels (**Figure 1A-B**). Using Bafilomycin A1 to disrupt lysosomal function, we found that loss of XPO1 activity led to increases in LC3 turnover, suggesting higher rates of autophagic flux (**Figure 1B-C**). Neuro2a cells treated with KPT-330 displayed enhanced lysosomal degradation of autophagy proteins LC3, SQSTM1, GABARAP as well as the nucleolar protein and ribosomal assembly modulator Fibrillarin (FBL) (**Figure 1D**) [14]. Transcriptionally, some lysosomal gene targets of the major autophagy transcription factor TFEB were induced suggesting improvements with lysosomal activity (**Figure 1E**), as previously shown in HeLa cells [8]. Altogether, our data suggested that KPT-330 could possibly improve proteostasis and amyloid clearance in mice neurons by stimulating the autophagy-lysosome pathway.

**Figure 1.**
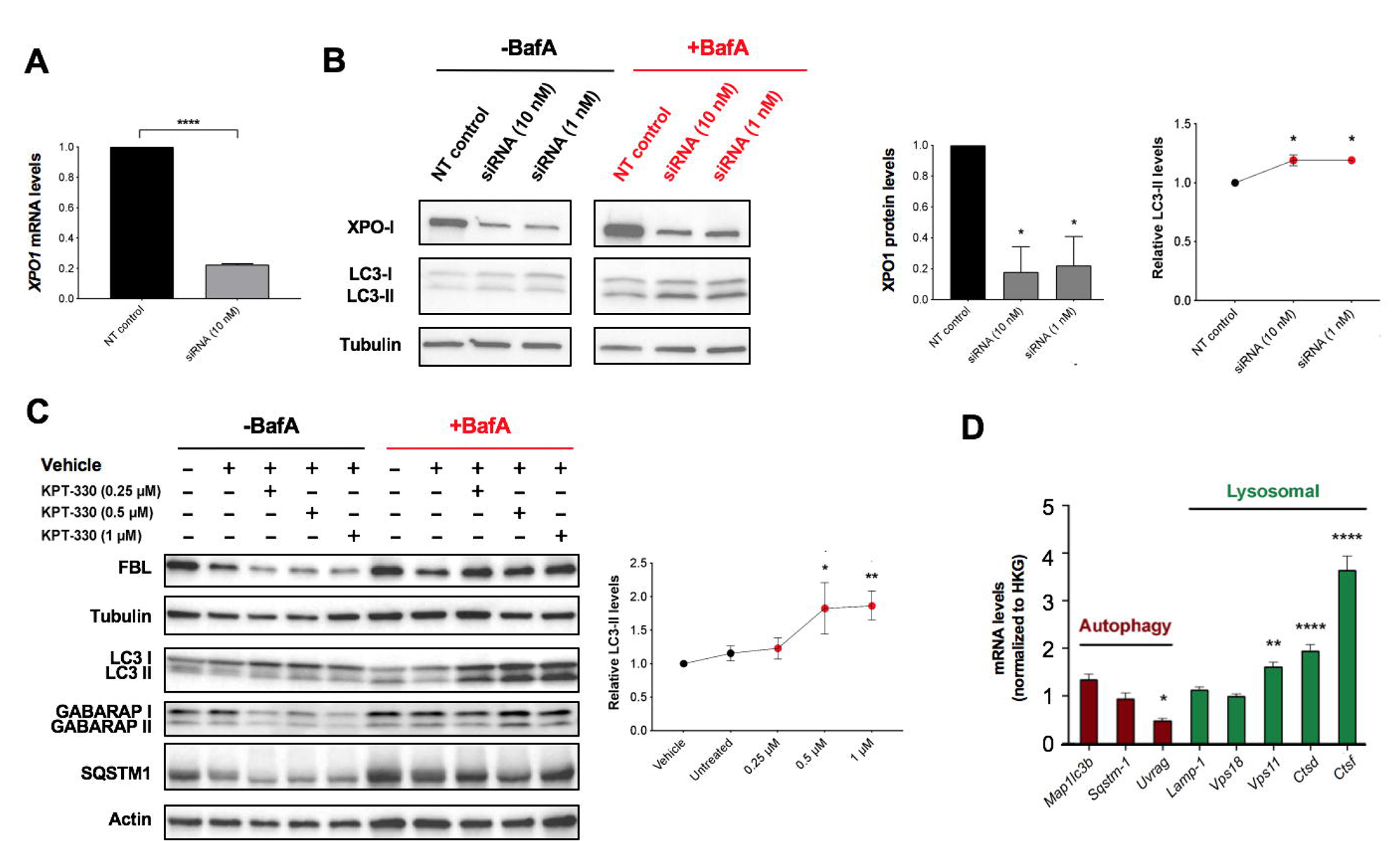
KPT-330 treatment increases autophagic degradation in Neuro2a cells. **A**. *XP01* silencing efficiency was measure by qPCR (n=3). **B**. Proteins in control and silenced Neuro2a cells were immunoblotted and levels of XP01 and LC3 forms (with Bafilomycin A1) were quantified (n=3, *ANOVA, *P<0*.*05* vs control). C. Neuro2a cells incubated with different concentrations of KPT-330 for 48 hours followed by a treatment with BafilomycinA1 for 6 hours and proteins were resolved by immunoblotting. LC3 forms were quantified (n=3, *ANOVA*, *: *P<0*.*05*, **: *P<0*.*01)*. D. Expression of autophagy and lysosomal genes (normalized to housekeeping genes, HKG) in cells treated with vehicle or KPT-330 (1uM) were quantified by qPCR (n=3, *: *P<0*.*05*, **: *P<0*.*01*, ****: *P<0*.*0001, t-test)*.

### Chronic KPT-330 treatment improves cognition in 5XFAD Alzheimer mice

As KPT-330 is expected to cross the blood brain barrier, we predicted that feeding KPT-330, an orally bioavailable compound [16], would lead to bioaccumulation in the brain. We found that KPT-330 readily accumulated in the blood and liver as well as the brain of C57BL/6J mice (**Figure S1A-C**). To test the effects of XPO1 inhibition on Alzheimer’s disease onset, we fed KPT-330 (low dose 2.6mg/kg/wk or high dose 45mg/kg/wk) (**Figure S1C)** [17] in C57BL/6J mice and the Alzheimer’s disease 5XFAD mice (hemizygous) [18] from 4 to 12 months of age. The rationale for the lower dose was to evaluate the impact of modest inhibition of XPO1, which could mimic conditions observed in long-lived nematodes that display partial reductions in XPO1 expression [13]. Chronic feeding of KPT-330 had minimal effect on weight (**Figure S1D**) and no discernable tolerability issue. We tested cognition after 6 months of age where amyloid deposition is established [19] via two complementary cognitive evaluations, the novel object recognition (NOR) for recognition memory [20] and spontaneous alternation in Y maze for spatial learning and memory. NOR testing revealed that KPT-330 did not improve novel object recognition (**Figures 2A-B, S2A-C**) [19]. However, our results showed that a high dose KPT-330 significantly increased spontaneous alternations in 5xFAD but not in WT mice suggesting an improvement in spatial memory (**Figure 2C-D**). Both NOR and Y maze tests showed tendencies for reduced overall distance traveled in 5XFAD mice treated with the high dose KPT-330 (**Figure 2B and 2D**), suggesting that these generally hyperactive animals [21] may be less active with KPT-330 treatment. Taken together, KPT-330 treatment resulted in functional improvements on spatial but not recognition memory, suggesting positive effects on XPO1 inhibition on spatial memory deficits in an Alzheimer’s disease model that may arise from improved brain proteostasis.

**Figure 2.**
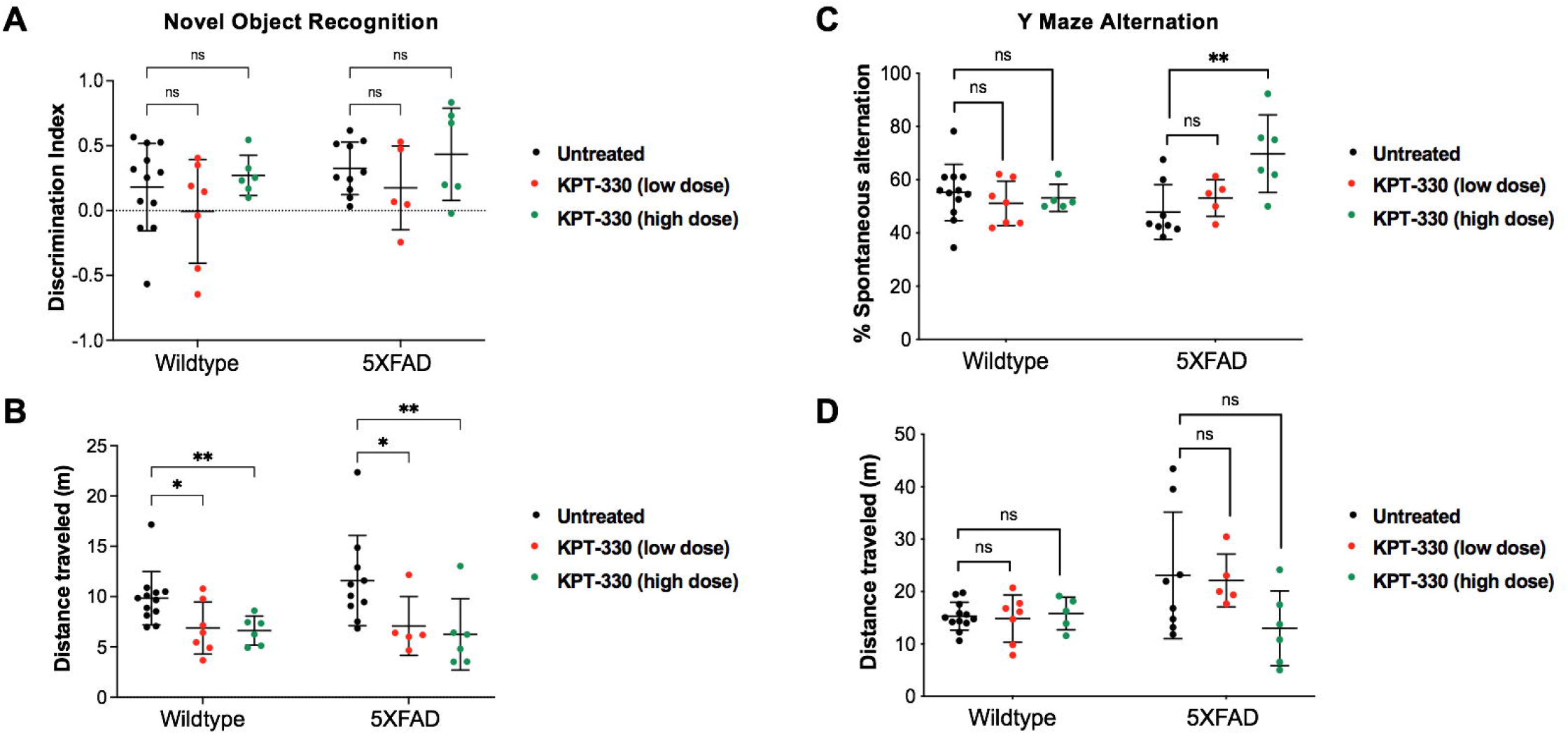
KPT-330 treatment improves cognition in SXFAD mice performing the spontaneous alternation Y maze test. **A**. Wild-type and SXFAD mice fed control chow or chow containing low or high dose of KPT-330 were subjected to novel object recognition where discrimination and **B**. distance were measured. **C**. Spontaneous alternation and **D**. distance traveled in a Y maze were measured in 12 months-old wild-type and SXFAD mice treated with or without KPT-330 for 8 months. ns: not significant,*: *P<0*.*05, **: P<0*.*01, n=6-12, ANOVA*.

### KPT-330 treatment does not affect vascular amyloid deposition

In order to fully characterize amyloid deposition in 5XFAD brains treated with KPT-330, we opted to visualize and quantify hemisphere-wide distribution of vascular and parenchymal amyloid deposition and plaques using 3D immuno-labeling and imaging via light sheet fluorescence microscopy. As amyloid buildup in vasculature may be indicative of impaired clearance [22], we compared amyloid deposition between control and high dose KPT-330 treated animals as we observed positive cognitive effects from high dose treatment. In both groups, plaque deposition was apparent throughout the cortex, in particular in deeper cortical layers, including isocortex layer 5. Overall, we observed minimal changes in vascular amyloid accumulation between control and treated mice (**Figure 3A-C, Data S1, Videos S1-2**) suggesting that the impact of KPT-330 in 5XFAD mice may specifically reside in brain regions relevant for cognition.

**Figure 3.**
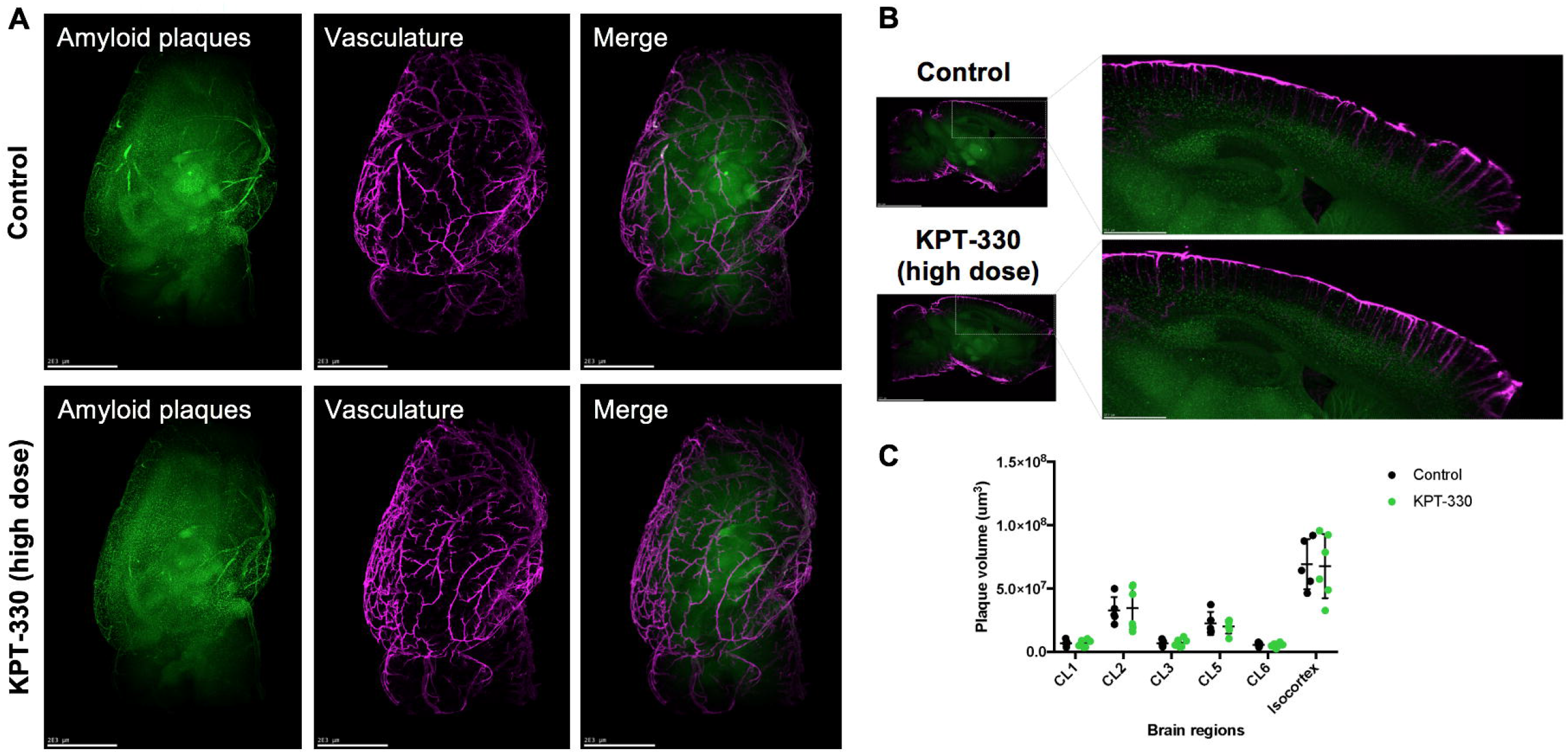
KPT-330 treatment does not affect vascular amyloid deposition. **A**. Representative images of SXFAD mice were treated with or without KPT-330 (high dose; 45mg/kg/week) for 8 months (Green: Congo Red, Purple: Anti-transgelin). B. Representative sagittal images of hemisphere staining of vasculature and amyloid plaques. C. Quantification of amyloid plaque volume in vasculature of 6 brain regions: CL1, 2, 3, 5 and 6, and isocortex (n=S-6 condition, *t-test)*. See Supplemental Videos S1-2 for 3D details.

### 5XFAD mice treated with KPT-330 differentially accumulate amyloid depots

By imaging and quantifying amyloid deposition in various brain regions, we unexpectedly observed an overall increase in amyloid plaques in the brain of 5XFAD treated with KPT-330, with deposition found in several auditory-related regions (**Figures 4A-C, S3, Data S1, Videos S1-2**). Accumulated amyloid plaque volume was significantly increased in several brain regions following treatment with KPT-330, including the magnocellular part of the lateral reticular nucleus, the caudal part of the spinal nucleus trigeminal, the nucleus of the trapezoid body, the ventral part of the medullary reticular nucleus, and the nucleus of the solitary tract. These unexpected amyloid accumulations were accompanied by higher overall ubiquitinated protein levels, suggesting potential regional proteostatic burden (**Figure S4A-B**). However, some brain regions had significantly lower amyloid levels (**Figure 4B**), including the dorsal penducular area, a site recently linked to anxiety-like behavior [23]. Further analysis in 12 relevant areas for amyloid deposition showed that thalamic regions displayed significant lower amyloid accumulation in 5XFAD animals treated with high dose KPT-330 (**Figure 4D**), suggesting that XPO1 inhibition cause differential amyloid deposition in the brain that protects decision-making regions. Finally, brain mRNA levels of autophagy genes LC3 and SQSTM1 were modestly increased by KPT-330 treatment (**Figure S4B-C**), in line with a transcriptional contribution to autophagy modulation by XPO1 (**Figure 1D**, [8, 13]). Taken together, our study shows that KPT-330 treatment impacts amyloid deposition in a region-specific manner and delays cognitive decline in a murine model of Alzheimer’s disease.

**Figure 4.**
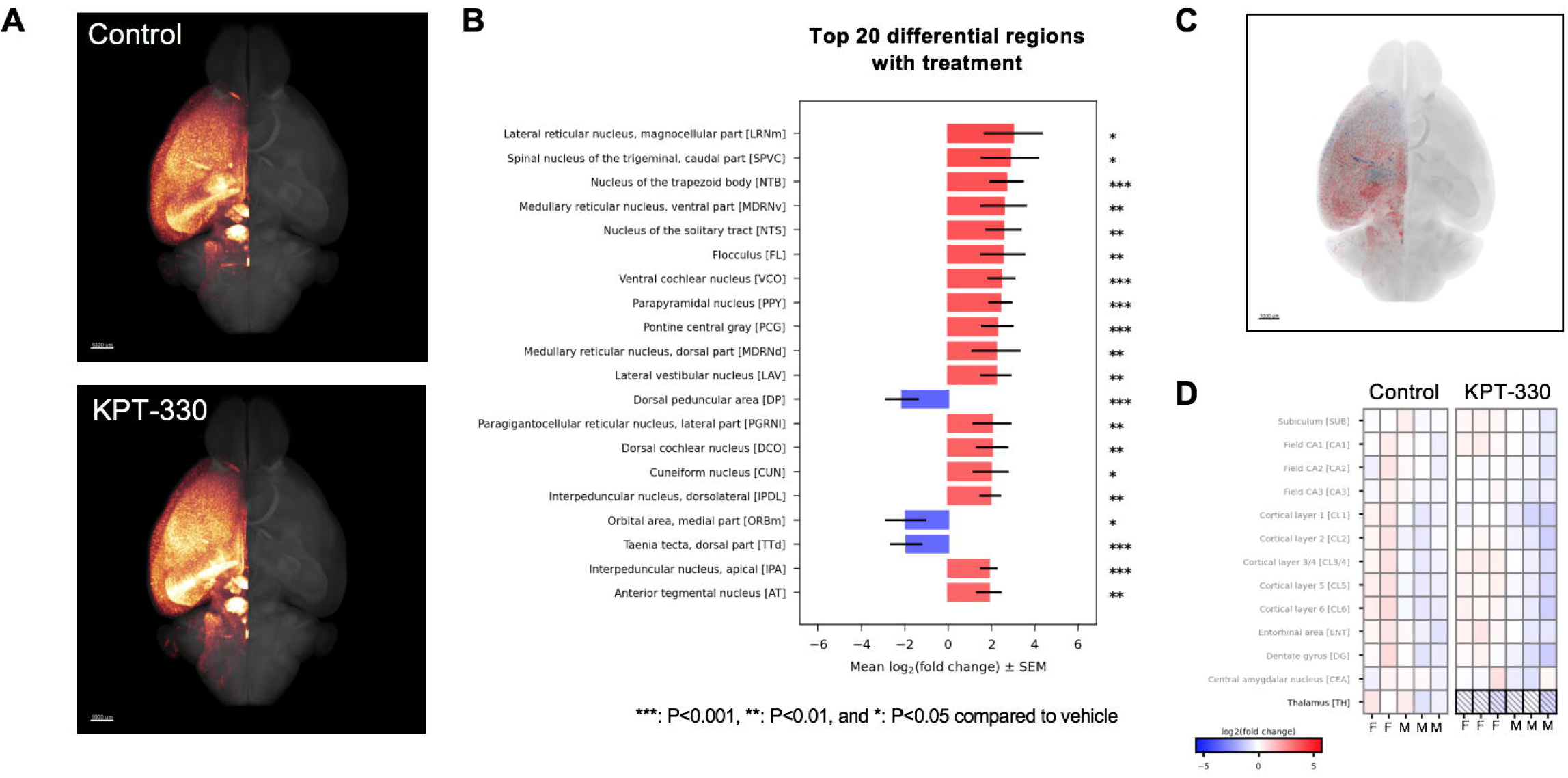
Amyloid differentially accumulates in the brain of 5XFAD mice treated with KPT-330. **A**. Representative group average of amyloid Congo Red staining (n=S-6/condition) in glow scale of brains from control and KPT-330 (high-dose; 45mg/kg/wk) treated 5XFAD mice. Scale bar = 1 mm. B. Top 20 significantly differential brain regions for amyloid deposition *(n=S-*6/condition, *t-test)* ***: P<0.001, **: P<0.01, and *: P<0.05 compared to control, visualized in **C. D**. Amyloid deposition in 12 selected brain regions reveal significant reduction in the thalamus P<0.05, n=S-6, *t-test*.

## DISCUSSION

Therapeutic development against Alzheimer’s disease remains challenging, and approaches to enhance neuronal proteostasis have yet to be approved clinically. Here, we evaluated the effect of XPO1 inhibition in murine cells and a murine model of Alzheimer’s disease and uncovered that loss of XPO1 leads to improvement in some cognitive domains. While amyloid plaque build-up is correlated with neuronal loss and progressive cognitive decline, loss of XPO1 activity generates differential amyloid depositions across regions of the brain that ultimately favor cognition-relevant areas.

Modulating autophagy and lysosomal biogenesis are potential approaches to enhance the clearance of proteotoxic proteins found in Alzheimer’s disease [24, 25]. Inhibition of XPO1 with KPT-330 leads to improvements in the autophagic turnover rate and lysosomal biogenesis in cells and showed promising effects in improving proteostasis and survival in models of neurodegenerative disease in small organisms [8]. Here, we find that acute inhibition of XPO1 in Neuro2a murine neuroblastoma cells also enhances autophagic flux. We anticipated that this effect may be recapitulated in the brains of Alzheimer’s disease mice treated with KPT-330. Y-maze analyses showed that spatial cognition is improved with KPT-330 at high dose, suggesting functional improvements across the brain and perhaps more specifically in regions important for cognition. Unexpectedly, the changes in amyloid deposition varied widely across different regions in the brain, indicating region-specific responses to XPO1 inhibition. In addition, KPT-330 had limited effects in vasculature amyloid deposition, suggesting that XPO1 inhibition affects plaque deposition directly in functional brain regions instead of via circulatory clearance. Our region-specific analyses (**Figure 4D**) agrees with the previously-described sex-specific differences in amyloid deposition in the 5XFAD mice model, whereas female mice brains tend to display higher levels of amyloid deposition [26]. Notably, significant increase in plaque accumulation in auditory regions was observed, a region of interest recently recognized in 5XFAD mice for amyloid deposition [27]. Conversely, thalamic regions accumulated significantly less amyloid plaques when mice were treated with KPT-330, highlighting protection in a region recognized for behavioral and cognitive processing [28, 29]. Thus, XPO1 inhibition in the brain of 5XFAD mice affects proteostasis differentially, indicating that modulation of nuclear protein export can lead to region-specific outcomes in amyloid accumulation. Overall, our study provides promising results on the benefits of XPO1 inhibition on cognition, but also reveals unexpected impact of KPT-330 on several brain regions. These spatially-specific effect of XPO1 inhibition in the brain warrant further studies as it may suggest cell type-specific responses in different brain regions from the loss of XPO1-mediated nuclear export, unraveling region-specific proteostatic flexibility that have ramifications in the development of Alzheimer’s disease.

## METHODS

### Immunoblotting in cells and tissues

Cells were washed with PBS and protein lysates were collected using RIPA buffer (50 mM Tris-HCl, 250 mM sucrose, 1 mM EDTA, and Roche protease inhibitor tablet, with 5% SDS), subjected to heat for 10 minutes. Tissues (10 mg) were homogenized with a pre-chilled Dounce homogenizer using RIPA buffer (50 mM Tris-HCl, 250 mM sucrose, 1 mM EDTA, and Roche protease inhibitor tablet). Tissue lysates were cleared by cold centrifugation. Lysate protein concentrations were quantified using the DC Protein Assay kit (BIO-RAD). Proteins were separated by SDS-PAGE on a 4-15% Tris-Glycine gel and transferred to nitrocellulose. The membrane was briefly stained with Ponceau S to confirm even transfer, rinsed with TBS-T until clear, blocked with 5% nonfat dry milk in TBS-T, and immunoblotted with anti-LC3 (Abcam, ab51520), anti-fibrillarin (Novus Biologicals, NB300-269), anti-SQSTM1 (Abcam, ab211324), anti-GABARAP (Abcam, ab109364), anti-Ubiquitin (ThermoFisher MAI-10035), anti-Tubulin (Abcam, ab6160) and anti-Actin (Milllipore, MAB1501R). Proteins were revealed using SuperSignal West Femto Maximum Sensitivity Substrate (ThermoFisher) and imaged on a ChemiDoc Imaging System (BIO-RAD). Densitometric analysis were conducted with ImageJ (NIH) and statistical analyses were performed using the Prism 7 software (GraphPad).

### mRNA extraction and analysis by qRT-PCR

Tissue and cell mRNA was extracted using Trizol Reagent and purified thereafter using Qiagen RNeasy kit. To perform qRT-PCR, cDNA was prepared as previously described [30]. qRT-PCR was performed using the QuantStudio 5 Real-Time PCR System (Applied Biosystems, ThermoFisher) and PowerTrack SYBR Green Master Mix (Applied Biosystems, ThermoFisher). The thermal cycling conditions were as follows: an initial denaturation at 95°C for 10 minutes, followed by 40 cycles of denaturation at 95°C for 15 seconds and annealing-extension at 60°C for 1 minute. Melt curve analysis was performed at the end of the reaction to verify the specificity of amplification. Data were analyzed using QuantStudio 5 software (Applied Biosystems, ThermoFisher). Expression of housekeeping genes (HKG) *B2m, Actb* and *Gapdh* were measured and genes of interest were normalized by the geometric mean of HKG as previously described [30]. Primer sequences can be found in **Supplemental Table S1**. The Prism 7 software (GraphPad) was used to conduct statistical analyses.

### Pharmacological analysis of KPT-330

KPT-330 (Adooq Bioscience) and fentanyl D5 (Millipore Sigma) super stock solutions were prepared in methanol at a concentration of 1 mg/ml. A working stock solution of KPT-330 was prepared each day from the super stock solutions at a concentration of 10 μg/ml and used to spike the calibrators. The LC/MS/MS system consisted of a Shimadzu SIL 20A HT autosampler, LC-20AD pumps, and an AB Sciex API 3200 tandem mass spectrometer with turbo ion spray. The LC analytical column was an ACE Excel C18-AR (75 x 3.0 mm, 3 micron, Mac-Mod Analytical) and was maintained at 25°C during the chromatographic runs using a Shimadzu CT-20A column oven. Mobile phase A contained 0.1% formic acid dissolved in water. Mobile phase B contained 0.1% formic acid dissolved in 100% HPLC grade acetonitrile. The flow rate of the mobile phase was 0.4 ml/min. KPT-330 and fentanyl D5 (the internal standard) were eluted with a gradient. The initial mobile phase was 25% B and at 1 minute after injection was ramped to 100% B. From 5.0 min to 8.0 min the mobile phase was maintained at 100% B and at 8.01 minutes was switched immediately back to 25% B and ran for 1.99 minutes to equilibrate the column before the next injection. The KPT-330 transition was detected in positive mode at 444/334.1 Da. The fentanyl D5 transition was detected at 342/188 Da.

In order to determine bioaccumulation of KPT-330, 4-months old mice received a bolus dose of KPT-330 (p.o.) and were then sacrificed at 60, 180, 360, 540, and 720 min post administration. Blood, brain, and liver were collected and assessed for KPT-330 concentration in each mouse. At each time point, n=3 male and n=3 female mice were included. KPT-330 was suspended in a solution of 5%Ethanol/0.6% Gibco® Pluronic® F-68/0.6% Polyvinylpyrrolidone (K30). KPT-330 was quantified in mouse whole blood. Calibrator samples were prepared daily by spiking blank blood to achieve final concentrations of 0, 1, 5, 10, 50, 100, and 500 ng/ml. Briefly, 0.1 mL of calibrator and unknown blood samples were mixed with 10 μL of 1 μg/mL fentanyl D5 and 1 ml of methanol. The samples were vortexed vigorously and then centrifuged at 13,000 *g* for 5 min at 25°C. The supernatants were transferred to a new 3ml tube and dried to residue under a gentle nitrogen stream. The residue was then redissolved in 100 μL of 50/50 mobile phase A/ mobile phase B, vortexing for 30 seconds. The samples were transferred to injection vials and 10 μL was injected into the LC/MS/MS. The ratios of KPT-330 peak areas to fentanyl D5 peak areas for each unknown sample were compared against a linear regression of the ratios obtained by the calibration samples to quantify KPT-330. The concentration of KPT-330 was expressed as ng/mL blood. Each brain and liver were weighed in a polypropylene tube and then a 10x volume of 75% methanol was added. Each sample was thoroughly homogenized. Calibrator samples were prepared daily by spiking a mixture of blank male and female control homogenate of each organ type to achieve final concentrations of 0, 1, 5, 10, 50, 100, and 500 ng/ml. The samples were prepared for injection and KPT-330 was detected as describe above. The concentration of KPT-330 was expressed as pg/mg of tissue.

### Mice maintenance, breeding and behavioral testing

Mice were raised on regular chow (NIH-31, Envigo) and 5XFAD hemizygous males were bred with B6J females (2 females per male ratio) and pups were raised and weaned by 21 days of age. Mice were fed control, low dose or high dose KPT-330 from 4 months of age onward. KPT-330 (Selleckchem) was added directly to food mixture and food pellet (Envigo). Wildtype and 5XFAD mice on control or KPT-330 chow were assessed for cognitive performance via the novel object recognition (NOR) and spontaneous alternation Y maze tests at 6 and 10 months of age respectively using the ANY-Maze video tracking and analysis software. Prior to testing, all animals were habituated to operator handling for 3 consecutive days at 1 min per animal during hours of the animals’ light cycle. On testing days, animals of different sexes were separately habituated in test rooms for 30 mins prior to the start of testing. All tests were conducted under low noise and lighting conditions (less than 30 lux) during hours of light cycle. All testing arenas and objects used were cleaned thoroughly with chlorhexidine and air dried to remove scent between animals. Animals which were tested were transferred to holding cages to reduce behavioral effects on untested littermates which may confound test readouts.

NOR assessment was adapted from [20]. Briefly, animals were first individually subjected to 30 minutes of habituation to an empty test arena approximately 30 x 30 x 30 cm, during which an open field test was concurrently performed to assess activity and anxiety levels (day 1). Animals were removed from the test arena and subjected to object familiarization 24 hours later (day 2). In this familiarization phase, animals were exposed to 2 identical objects placed at different corners of the test arena for 10 mins, during which object bias was determined by the following metric:

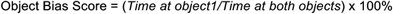

Animals exhibiting an object bias score below 20% or above 80% were excluded from subsequent analysis. In the test phase (day 3; 24 hours later), animals were returned to the test arena with one familiar object from the previous day and one novel object, both at the same positions as before, for 10 mins. The ability of the animals to distinguish novel from familiar objects was determined by two complementary measures, novelty preference (%) and discrimination index:

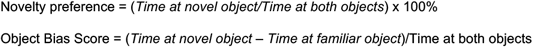

Preferential interaction with novel objects is suggestive of recognition for previously-encountered familiar object and is indicated with preference scores above 50%, whereas scores of 0-50% indicate no novel object preference. Similarly, positive and negative discrimination indexes indicate novel object and familiar object preferences respectively. Object interaction is scored if the animal was within 2 cm of it in the absence of object mounting.

The spontaneous alternation Y maze test was previously described in [31]. Briefly animals were placed into the middle of the starting arm and allowed to explore all arms of the maze for 8 minutes. Entry into an arm is scored when all four paws of the animals were inside the arm. For a spontaneous alternation to occur, the animal must enter a different arm of the maze consecutively and is calculated as follows:

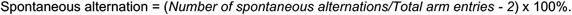

Increasing percentage of spontaneous alternations is suggestive of enhanced spatial memory.

### Brain imaging and analysis

Brains were dissected and fixed by perfusion. Intact hemispheres were stained using an adapted version of the previously published protocol for immunolabelling-enabled 3D imaging of solvent cleared organs (iDISCO) [32]. Beta-amyloid plaques were stained using Congo Red (15μM, #C6277, BioXtra, Sigma Aldrich) and the vasculature was immunostained with an anti-transgelin-targeting (SM22) antibody (1:1000, ab14106, Abcam) and donkey anti-rabbit (1:1000, 711-655-152, Jackson Immunoresearch). Following tissue clearing, the intact hemispheres were imaged using alight sheet fluorescence microscope (LSFM) (UltraMicropscope II, Miltenyi Biotec GmbH, Bergisch Gladbach, Germany) (Gubra). The hemispheres were positioned in a sagittal orientation within a dibenzyl ether-filled chamber for refractive index matching. Double-sided illumination was applied with an exposure time of 232 ms, capturing images at 10 μm intervals at 1.2x total magnification. Emission wavelengths of 500 nm (tissue autofluorescence), 560 nm (Congo Red), and 785 nm (SM22) were used for data acquisition. For each sample, hemisphere-wide plaque distribution was detected by Congo Red and the plaque volume was mapped to one of more than 800+ unique brain regions in a LSFM optimized reference atlas [33]. Plaques were divided into two populations depending on the distance to a blood vessel; >3 μm distance categorizes as parenchymal and <3 μm categorizes as vascular. Regions expressing significant regulation were presented via bar plots of plaque volume and group-wise statistics. All region names and abbreviations follow Allen’s Brain Atlas nomenclature. For each region, a negative binomial generalized linear model (GLM) was fitted to the quantitative data, and a subsequent Dunnett’s test was performed for multiple comparisons. Significantly regulated regions went through statistical validation, where the model deviance residuals were investigated to see if they aligned with the assumptions of Normality and homoscedasticity. Cook’s distance was evaluated for each data point to make sure no point is overly influential in the fitted model. If a region did not pass the validation, it was not considered significant. Notably, significance may be affected by the integrity of brain regions (i.e. dissection) and by abundance of plaque in the region (no lower threshold was set on plaque size and volume, meaning that few plaques in small regions can drive significance). Plaque volume was quantified in individual brain regions, further the vascular plaque volume was determined by layer-wise analysis in the isocortex. The effect of treatment between groups was assessed in 12 preselected regions by statistical analysis.

## ACKNOWLEDGEMENTS

We are grateful for the support in animal breeding and care provided by Rebecca Poncin, Lauren Souza and Lara Helwig at the Brown Animal Care Facility. The study in mice was performed according to the IACUC protocol #20-06-0003 and was carried out at Brown University (AAALAC accredited). We would like to thank Marisa Lopez-Cruzan, Martin Javors and Peter Hornsby at UT San Antonio – Nathan Shock Center for their help in coordinating the pharmacological analyses. We would like to thank Lea Lydolph Larsen (Gubra) for coordinating brain staining, 3D imaging and analysis. The transgenic mouse strain, B6.Cg-Tg(APPSwFlLon,PSEN1*M146L*L286V)6799Vas/Mmjax,RRID:MMRRC_034848-JAX, was obtained from the Mutant Mouse Resource and Research Center (MMRRC) at The Jackson Laboratory, an NIH-funded strain repository, and was donated to the MMRRC by Robert Vassar, Ph.D., Northwestern University.

## AUTHORS CONTRIBUTIONS

SQW performed the behavioral study and analysis, mRNA and protein analyses. AO conducted mRNA and protein analyses. AM oversaw mice maintenance, and RT and AA performed feeding and weighing throughout the study. ANM, JJS and BCG coordinated and conducted the pharmacological analysis. AAA oversaw the mice behavioral study and analysis. LRL managed the study and wrote the manuscript. All co-authors edited the manuscript.

