## Supplemental Material for "Spatial memory in Alzheimer’s disease 5XFAD mice is enhanced by XPO1 inhibitor KPT-330"

Figure S1.

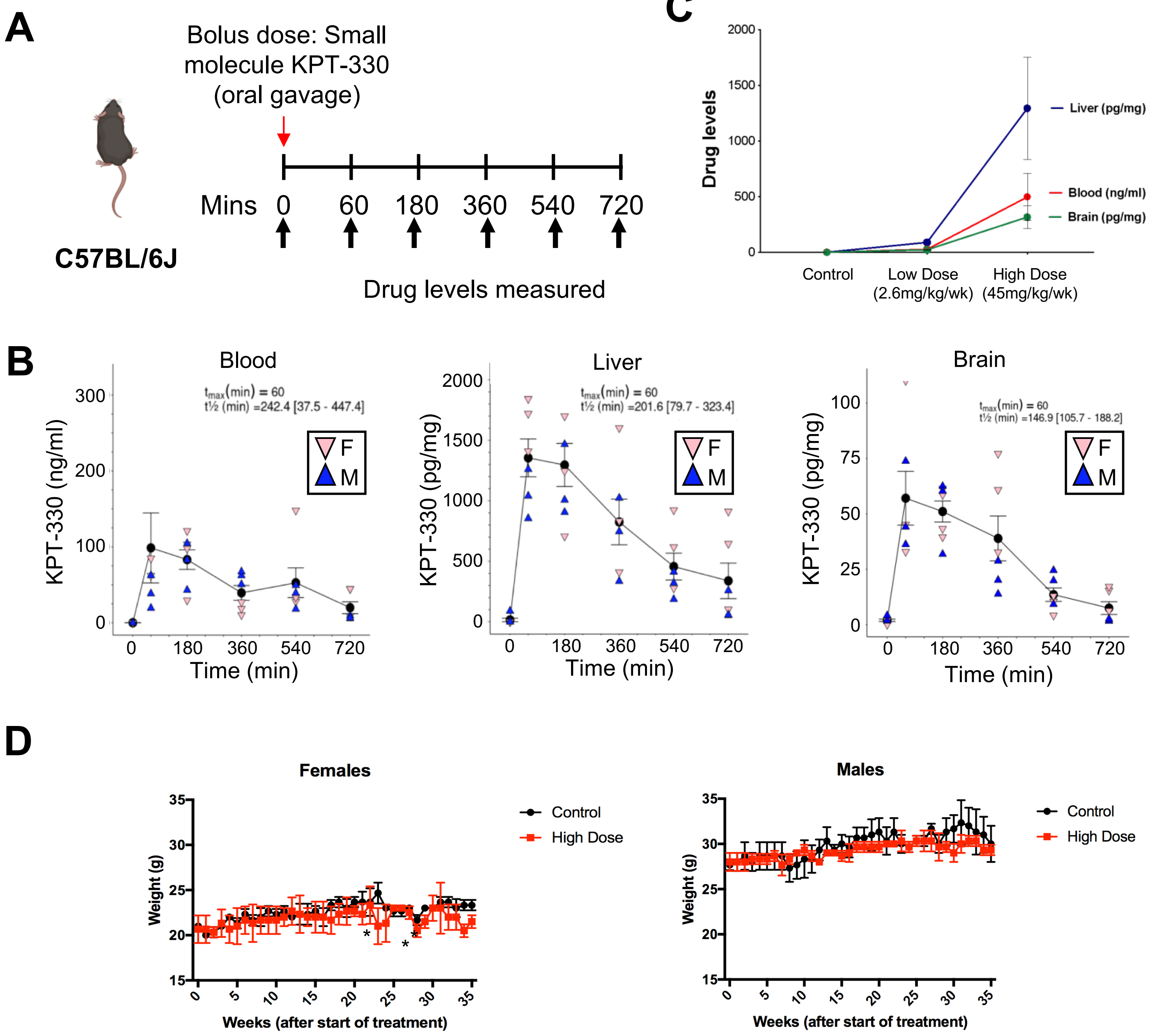


**Figure S1. Pharmacological and bioaccumulation analysis of KPT-330. A.** C57/B6 mice received a bolus dose of 7.5mg/kg KPT-330 (p.o.) and were then sacrificed at 60, 180, 360, 540, and 720 min post administration. **B.** Blood, liver and brain were collected and assessed for KPT-330 concentration in each mouse. At each time point, 4-month old male (n=3) and female (n=3) mice were included. **C.** Blood, liver, and brain levels of KPT-330 were also analyzed in 4-month old mice fed chow, or chow with low or high dose of KPT-330 for two weeks. **D.** Mice weight was measured for the duration of the treatment (45mg/kg/wk) from 4-months to 12 months of age (n=3 per sex per treatment) **: $P<0.05$, t-test.*

Figure S2.

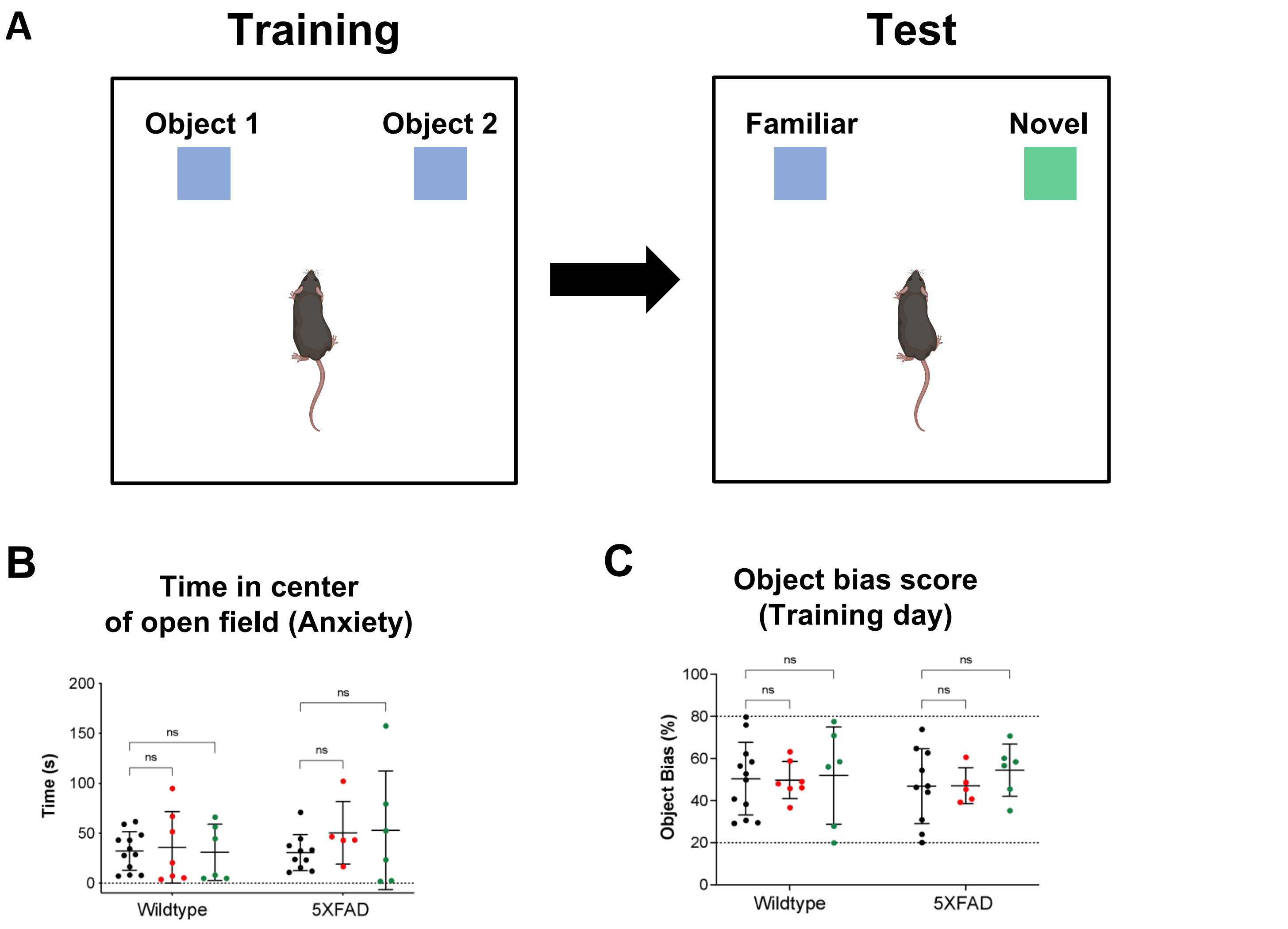


**Figure S2. KPT-330 treatment does not improve novel object recognition (NOR) in 5XFAD mice. A.** Four-month old wildtype and 5XFAD mice were fed with either chow or chow containing high (45 mg/kg/wk) or low (2.6 mg/kg/wk) dose of the XPO1 inhibitor, KPT-330, for 2 months and assessed for effects on recognition memory using the NOR test at 6 months of age following prior habituation to the test arena and object familiarization (see methods for details). **B.** Time in center of open field and **C.** object bias score during training day were measured. ns: not significant, n=6-12, *ANOVA.*

Figure S3.

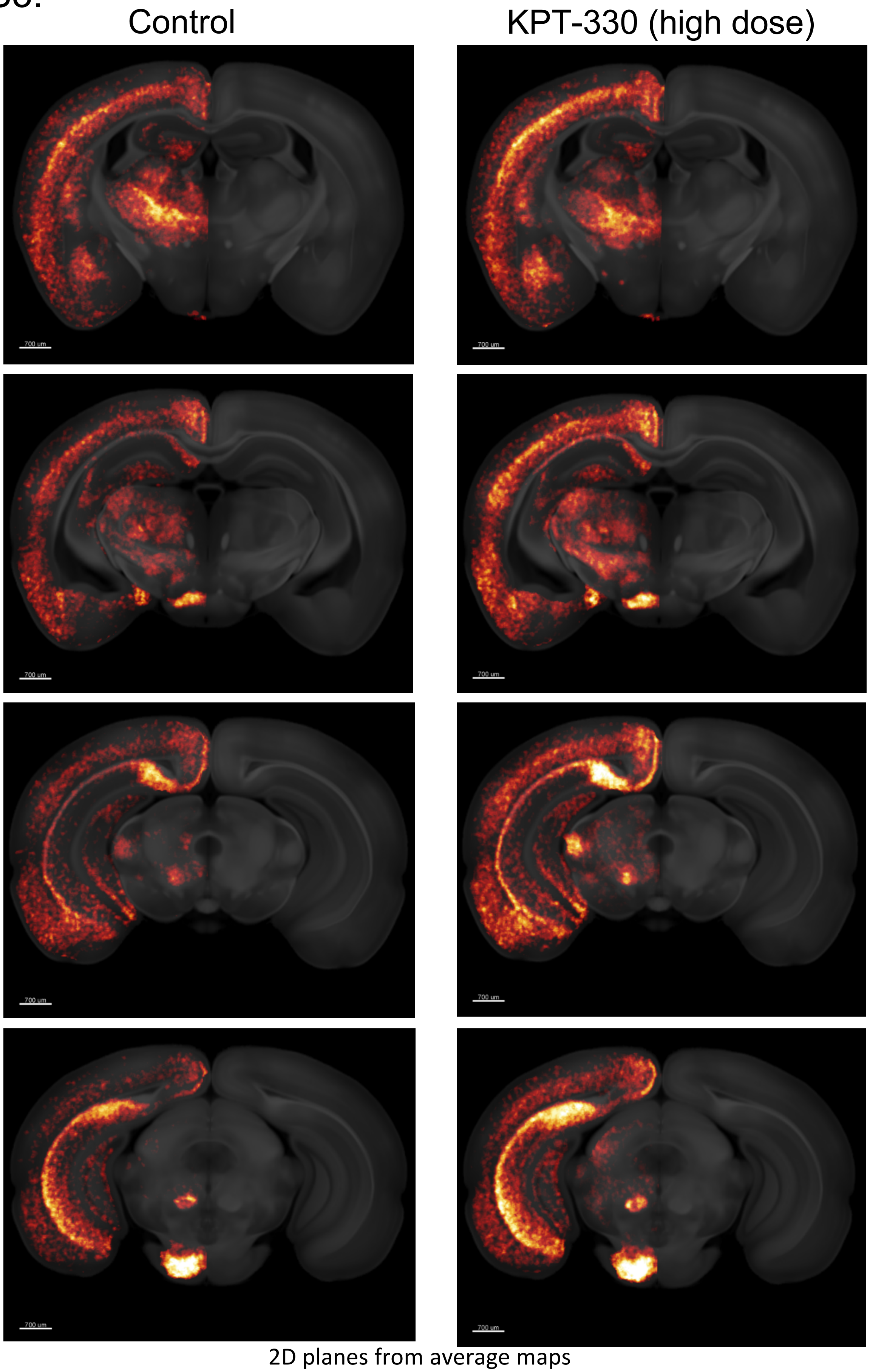


**Figure S3. Differential amyloid accumulation in 5XFAD mice brains.** Representative 2D images at selected coronal planes showing group average of plaque volume in response to KPT-330 treatment and with control shown in glow scale. Scale bar 700 µm.

Figure S4.

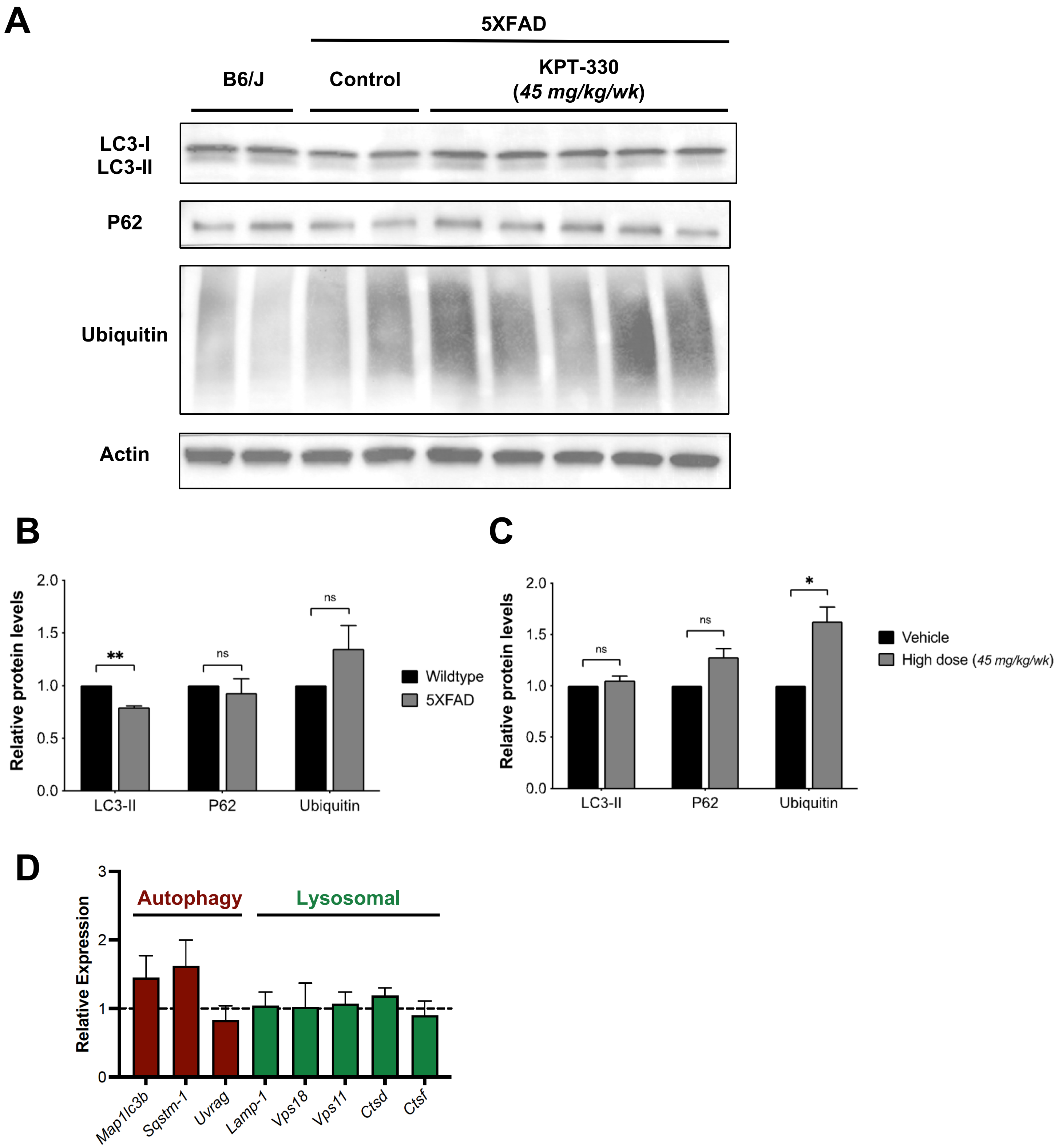


**Figure S4. Protein analysis in brains of 5XFAD mice. A.** Immunoblotting of LC3, p62 (SQSTM1) and ubiquitin in brains of 9-months old B6/J and transgenic 5XFAD mice (with or without KPT-330 from 4 months of age). **B.** Relative protein levels was compared between B6/J (wild-type) and 5XFAD (**B**) and between control (vehicle) and high dose KPT-330 (**C**). Values expressed as mean of *n* = 2 ± SEM; *: *P<0.05*, **: *P<0.01.* **D.** mRNA levels of autophagy and lysosomal genes in the brain of 9-month old 5XFAD animals and treated with KPT-330 compared to control (normalized at 1), n=3, *t-test*.

### Table S1.

| Gene name | Sequence 5' -3' |
|---|---|
| *Gapdh* Forward | CATCACTGCCACCCAGAAGACTG |
| *Gapdh* Reverse | ATGCCAGTGAGCTTCCCGTTCAG |
| *Actb* Forward | GTGACGTTGACATCCGTAAAGA |
| *Actb* Reverse | GCCGGACTCATCGTACTCC |
| *B2m* Forward | ACCCGCCTCACATTGAAATCC |
| *B2m* Reverse | GGCGTATGTATCAGTCTCAGTG |
| *Map1lc3b* Forward | CGGACTGAGACACACACAAGG |
| *Map1lc3b* Reverse | TCGCTCTATAATCACTGGGATCT |
| *Sqstm1* Forward | GGCCTATCTTCTGGGCAAGG |
| *Sqstm1* Reverse | CCGGCACTCCTTCTTCTCTTT |
| *Uvrag* Forward | ATCCGACGTGGAGGAGTCTT |
| *Uvrag* Reverse | TGCGTTTGGATGACCCTTGT |
| *Lamp 1* Forward | TGCTCCGGGATGCCACTAT |
| *Lamp 1* Reverse | CTTGTTGTCCTTTTTCAGGTAGGTG |
| *Vps18* Forward | CGCCTGCGTCCATGTCTATAA |
| *Vps18* Reverse | GGCAGCACATCCTCGATCTT |
| *Vps11* Forward | CTGGAAAAGAGAGACGGTGGCAATC |
| *Vps11* Reverse | GAAAGGCCAGCCCAGTAACG |
| *Ctsd* Forward | CTGTGGGTCCCCTCCATTCATTG |
| *Ctsd* Reverse | GGTAGCCCATGCCCAAGATG |
| *Ctsf* Forward | CACAGCTCAGTATGGGATCACC |
| *Ctsf* Reverse | TTGGCTGGACTCATCTTCCTGC |

**Table S1. Primer used in the study**

Data S1.

See Excel File

Video S1.

See Video File

Video S2.

See Video File
